# EpiESM-GA: Resource-Efficient Protein Foundation Model Features for Equitable B-Cell Epitope Prediction

**DOI:** 10.64898/2026.06.22.733745

**Authors:** Purnima Gautam, Pralay Mitra

**Affiliations:** Department of Computer Science & Engineering Indian Institute of Technology Kharagpur Kharagpur, India

**Keywords:** B-cell epitope prediction, machine learning, protein foundation models, ESM-2 embeddings, evolutionary feature selection

## Abstract

Prediction of B-cell epitopes can assist in reducing costly wet-lab screening in vaccine design, diagnostics, and antibody discovery. However, current predictors often suffer from noisy labels, weak generalization, and structure-dependent workflows. Here we present EpiESM-GA, an efficient sequence-only pipeline for linear B-cell epitope prediction. Positive and negative peptide examples are collected from IEDB, which provides experimentally tested epitopes and distinguishes positive and negative epitope records based on assay evidence(Vita et al., 2019). Each peptide is encoded with a *frozen* ESM-2 protein language model: a bidirectional transformer producing amino acid embeddings for downstream structure and function tasks (Lin et al., 2023). Mean-pooled embeddings are further compressed into a compact 420-feature representation with a genetic algorithm and classified with lightweight Random Forest, XGBoost, or MLP heads. This avoids foundation-model fine-tuning, reduces the number of trainable parameters, improves interpretability, and enables low-resource deployment. On an IEDB-derived benchmark, EpiESM-GA attains **0.880**± **0.004** AUC-ROC, **0.852**± **0.005** PR-AUC, **82.0** ± **0.6%** accuracy, **0.79** ± **0.01** F1, and **0.74**± **0.01** MCC, outperforming dense ESM-2 features and baselines LBCE-XGB, EpitopeVec, and BepiPred-2.0 (mean± std over five independent random seeds). The framework shows how frozen protein foundation models can enable pandemic preparedness, peptide vaccine prioritization, diagnostic antigen screening, and equitable computational immunology.

## 1 Introduction

Proteins are sequential, like human language. Amino acids are discrete tokens, protein families have grammar-like regularities, and functional constraints leave statistical signatures in evolutionary sequence space. Protein language models (PLMs) exploit this structure by training transformer encoders on large corpora of proteins, allowing structural, functional, and evolutionary signals to emerge from the primary sequence alone (Rives et al., 2021; Lin et al., 2023). This makes PLMs attractive for biomedical social-good problems where wet-lab screening is expensive, slow, and unevenly available across institutions and countries.

One application is the prediction of B-cell epitopes. B-cell epitopes are parts of antigens that are recognized by antibodies and are the target of peptide vaccine design, immunodiagnostics, and therapeutic antibody engineering (Potocnakova et al., 2016; Caoili, 2022). Peptide arrays, crystallography, and immunoassays are still the gold standard for experimental mapping but require specialized infrastructure, reagents, and time. Thus, a reliable sequence-only predictor can be used as a triage tool—not a substitute for experiments but to narrow the candidate search space and to identify high-confidence peptides, especially in the context of emerging-disease outbreaks.

Computational epitope prediction has evolved from early propensity-scale methods to supervised machine-learning models based on amino acid composition, peptide kernels, and ensemble classifiers. More recently, performance of PLM-based methods has been improved: EpitopeVec (Bahai et al., 2021) applies a support vector classifier on protein embeddings, LBCE-XGB (Zhu et al., 2023) combines BERT-style embeddings with XGBoost, and BepiPred-3.0 (Clifford et al., 2022) uses ESM-1b representations to predict linear and conformational epitopes. However, there are still key challenges: heterogeneous assay-derived labels, limited cross-dataset robustness, and the cost of pipelines requiring full model fine-tuning or structural information. These constraints are especially severe for small labs and public health groups in low-resource settings.

Responsible foundation-model deployment should be accurate, efficient, auditable, and usable without industrial-scale compute (Strubell et al., 2019; Schwartz et al., 2020; Patterson et al., 2021). We introduce EPIESM-GA, a pipeline that curates IEDB peptide records, encodes sequences with a frozen ESM-2 model, selects a compact 420-dimensional embedding subset using a genetic algorithm wrapper, and trains lightweight downstream classifiers.

Our contributions include: (i) a no-fine-tuning PLM pipeline for linear B-cell epitope prediction; (ii) GA-based feature selection outperforming PCA and random compression at equal dimensionality; (iii) ablations over handcrafted, full ESM-2, compressed, and GA-selected features; and (iv) a deployment-oriented discussion, including pandemic preparedness, vaccine prioritization, diagnostic antigen screening, calibration, cross-validation, and controlled baseline evaluation.

## 2 Related Work

### 2.1 Prediction of B-cell epitopes

Early B-cell epitope predictors used manually constructed propensity scales based on hydrophilicity, flexibility and semi-empirical antigenicity (Parker et al., 1986; Kolaskar and Tongaonkar, 1990). Later supervised approaches introduced neural networks, string kernels, support vector machines and ensemble feature design as shown in ABCPred, BCPred, SVMTriP, LBtope and iLBE (Saha and Raghava, 2006; El-Manzalawy et al., 2008; Yao et al., 2012; Singh et al., 2013; Hasan and Kurata, 2020). BepiPred-2.0 improved sequence-based prediction using a random forest trained on structure-derived epitopes (Jespersen et al., 2017), and deep models trained on IEDB-scale data further advanced linear epitope prediction (Liu et al., 2020). More recently, protein language model (PLM) embeddings are increasingly adopted: EpitopeVec uses protein sequence embeddings (Bahai et al., 2021), LBCE-XGB combines BERT-style embeddings with XGBoost (Zhu et al., 2023), and BepiPred-3.0 employs ESM-1b residue-level representations with a trained linear head for both linear and conformational epitope prediction (Clifford et al., 2022; Gautam and Mitra, 2025). EPIESM-GA also follows this PLM-based direction, but focuses on compact feature selection and low-resource deployment rather than end-to-end training.

### 2.2 Protein language models

Recent PLMs are built on transformer architectures (Vaswani et al., 2017) and BERT-style masked language modelling (Devlin et al., 2019). ESM and ProtTrans learn evolutionary and structural constraints from large corpora of proteins (Rives et al., 2021; Lin et al., 2023). AlphaFold (Jumper et al., 2021), ESMFold and ESM-3 further illustrate the ability of learned protein representations to facilitate structural and multimodal sequence–structure–function modelling (Hayes et al., 2024).

### 2.3 Responsible and efficient foundation models

For efficient deployment of foundation models, pruning, distillation, quantization, or low-rank adaptation (Han et al., 2016; Hinton et al., 2015; Sanh et al., 2019; Hu et al., 2022; Dettmers et al., 2023) are often adopted. For biomedical sequence tasks, freezing the encoder and caching embeddings to train small task-specific models provides a complementary route aligned with Green AI, where efficiency and access matter alongside accuracy (Schwartz et al., 2020). Unlike model compression, genetic-algorithm wrapper selection selects downstream feature indices instead of modifying the PLM, which makes peptide screening more feasible for labs with limited GPU access.

## 3 Data and Experimental Setup

### 3.1 Problem definition

Given an amino acid peptide *x* = (*a*_1_, …, *a*_*L*_), where *a*_*i*_ is one of the 20 canonical residues and *L* is the peptide length, we pose linear B-cell epitope prediction as binary classification

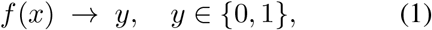

where *y* = 1 denotes a positive linear B-cell epitope and *y* = 0 denotes a non-epitope. The predicted score *p*(*y* = 1 |*x*) is used to score candidate peptides for downstream experimental validation.

### 3.2 IEDB curation and sequence-identity splits

We curate peptide records from the Immune Epitope Database (IEDB), a manually curated resource of experimentally characterized immune epitopes from antibody, T-cell, and MHC-binding assays (Vita et al., 2019; Peters et al., 2012). Positive samples are linear B-cell peptides with at least one positive assay outcome. Negative samples are peptides reported as non-reactive across all IEDB assay records in which they appear, giving experimentally grounded negatives rather than randomly sampled unannotated antigen regions.

#### Cleaning

We remove duplicate sequences, entries with missing or conflicting labels, and peptides containing non-canonical or ambiguous residues (B, J, O, U, X, Z). We retain peptides of length 8-50 amino acids, covering more than 99% of linear B-cell epitopes in IEDB and remaining below the ESM-2 token limit of 1,022 residues. The cleaned dataset has median length 15 residues, with interquartile range 12-20.

#### Sequence-identity splitting

To reduce homology leakage, a known source of inflated epitope benchmark performance (Jespersen et al., 2017; Clifford et al., 2022), we cluster peptides at 90% sequence identity using CD-HIT (Fu et al., 2012). Entire clusters are assigned to one split only.

#### Class balance and splits

Matched negative sampling balances each split. Validation is used for GA fitness and early stopping; the test set is reserved for final reporting.

## 4 EpiESM-GA Methodology

### 4.1 Pipeline overview

Figure 1 summarizes EPIESM-GA. Peptide sequences are tokenized, encoded by a frozen ESM-2 model, mean-pooled into 1280-dimensional sequence embeddings, compressed to 420 task-relevant dimensions by a genetic-algorithm selec-tor, and classified using lightweight downstream models.

**Figure 1.**
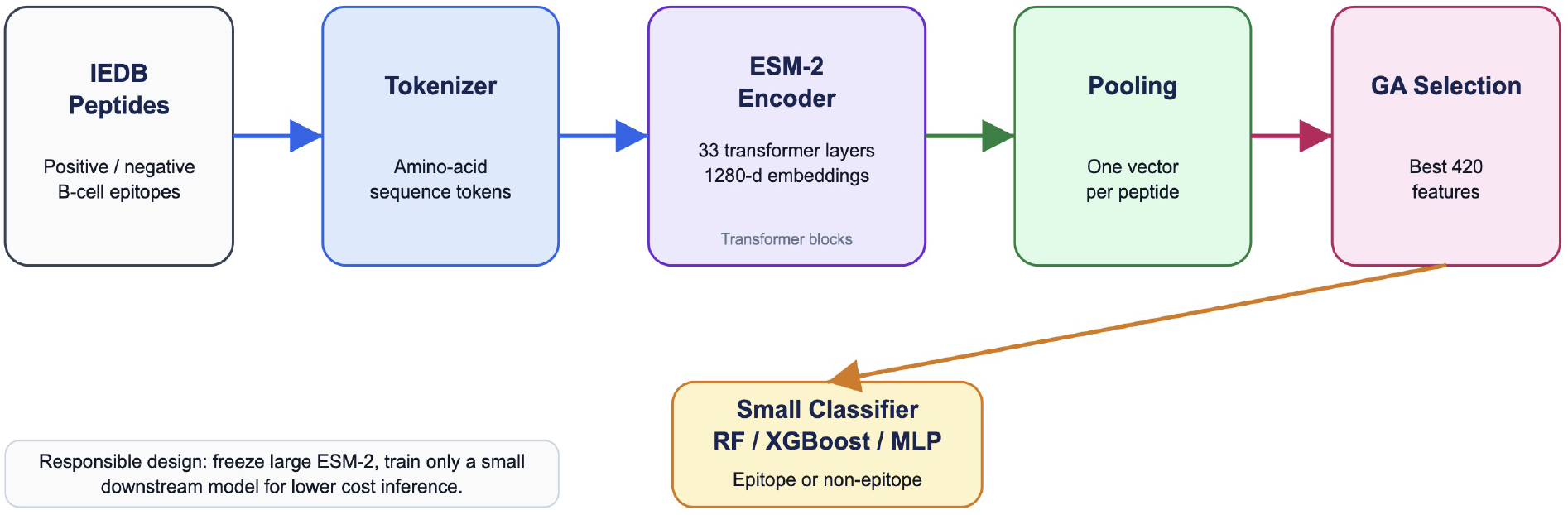
EpiESM-GA pipeline. IEDB peptides are tokenized, embedded with a *frozen* ESM-2 encoder (33 transformer layers, 1280-dim hidden states), mean-pooled to one sequence vector per peptide, reduced to 420 dimensions by a genetic-algorithm mask, and classified by a lightweight downstream model.

**Figure 2.**
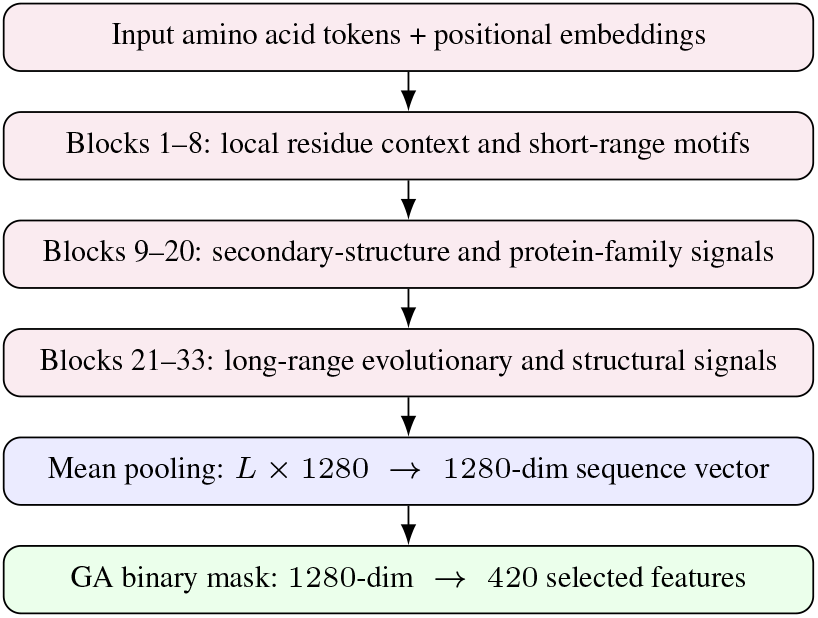
ESM-2 feature creation in EpiESM-GA. The encoder is fully frozen. Layer-group characterisation follows Lin et al. (2023).

### 4.2 ESM-2 feature creation

ESM-2 is a bidirectional transformer encoder trained with masked language modelling over 250 million protein sequences (Lin et al., 2023). For a peptide *x* = (*a*_1_, …, *a*_*L*_), token and positional embeddings define

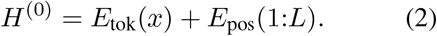

Each transformer block applies multi-head self-attention, residual connections, layer normalization, and a feed-forward network:

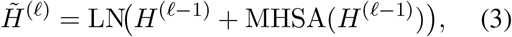

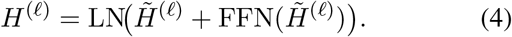

We use the 650M-parameter ESM-2 variant with 33 layers and 1280-dimensional hidden states. All encoder parameters are frozen, and only the final-layer residue embeddings *H*^(33)^ ∈ ℝ^*L×*1280^ are extracted. A smaller 150M-parameter variant achieved AUC within approximately 0.01 in preliminary tests, suggesting a useful future trade-off for highly resource-constrained deployment.

A sequence-level representation is obtained by mean pooling:

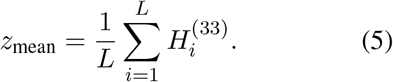

Mean pooling is stable for short peptide fragments, which dominate linear B-cell epitope datasets. We leave max pooling and biochemical feature concatenation for future work to keep the core pipeline dependency-light.

### 4.3 Evolutionary feature selection

Although 1280-dimensional PLM embeddings are expressive, many dimensions are redundant for epitope classification. Following genetic-algorithm wrapper selection for peptide descriptors (Angaitkar et al., 2023), we represent each candidate subset as a binary mask *m* ∈ {0, 1}^1280^ and compute masked features *z*_*m*_ = *z* ⊙ *m*.

#### Fitness function

The GA maximizes validation AUC while penalizing feature count:

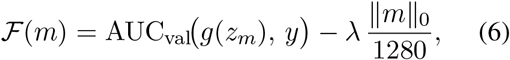

where *g* is an XGBoost classifier trained on masked training features and evaluated only on validation data, with *λ* = 0.01 following Angaitkar et al. (2023). The held-out test set is used once after feature selection and classifier decisions are finalized.

#### GA configuration and leakage control

We run the GA with 100 masks for 50 generations, initializing each mask with approximately 420 active dimensions. Tournament selection uses size 5, with uniform crossover rate 0.7, bit-flip mutation rate 0.01, and top-5 elitism. A repair step restores ∥*m*∥ _0_ ≈ 420 after variation. The validation split is used only for GA fitness and early stopping; classifier settings are fixed before evolution.

### 4.4 Downstream classifiers

We evaluate four lightweight classifiers on the GA-selected 420-dimensional features. Random Forest provides an interpretable ensemble baseline and feature-importance estimates. XGBoost models nonlinear interactions efficiently (Chen and Guestrin, 2016). A small multilayer perceptron with two hidden layers tests whether a neural head can extract additional signal without PLM fine-tuning. Logistic regression serves as a linear and calibration-friendly baseline. All classifiers are trained only on training-split features; validation labels are not used for classifier weight optimization.

#### Algorithm 1

Evolutionary selection of ESM-2 fea-ture dimensions

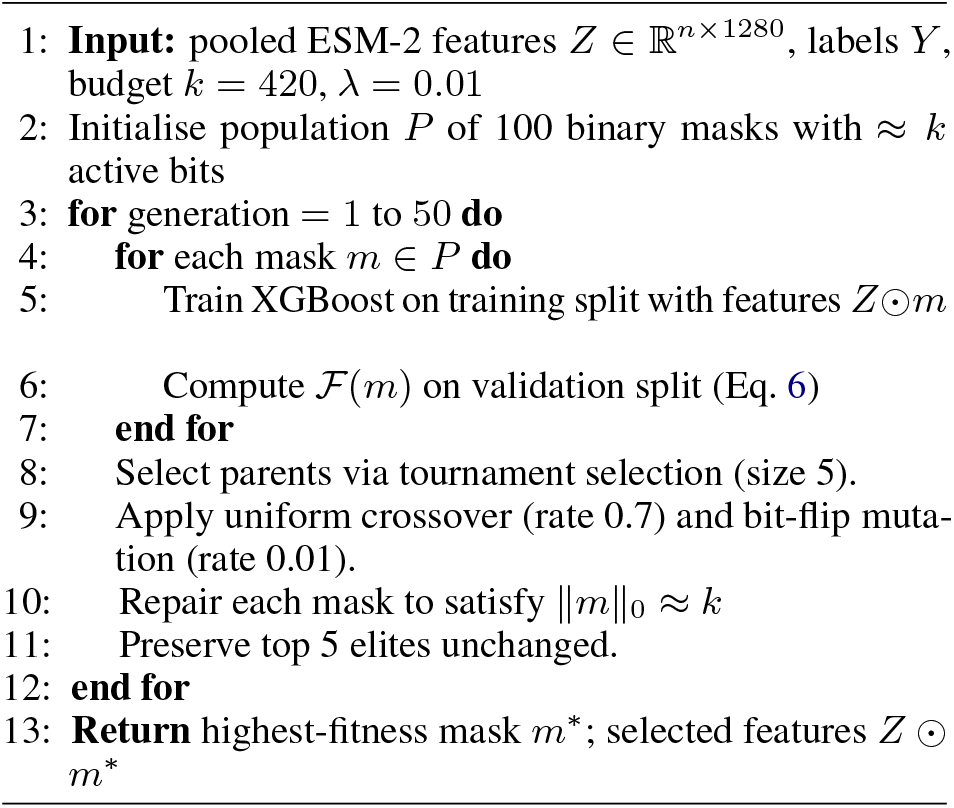

## 5 Experiments

### 5.1 Baselines

We compare EPIESM-GA against six baseline groups: (i) handcrafted sequence features, including amino acid composition and dipeptide composition; (ii) BepiPred-2.0, a random forest trained on structure-derived epitopes (Jespersen et al., 2017); (iii) EpitopeVec, which uses PLM embeddings with a support vector classifier (Bahai et al., 2021);(iv) LBCE-XGB, which combines BERT-style embeddings with XGBoost (Zhu et al., 2023); (v) BepiPred-3.0, the most relevant PLM-based baseline for linear epitope prediction (Clifford et al., 2022); and (vi) internal ESM-2 baselines using either all 1280 pooled dimensions or a random 420-dimensional subset. The random-subset baseline tests whether the GA provides discriminative selection rather than merely reducing dimensionality. Results from published systems are indicative because their original test sets differ from ours; the ESM-2 baselines are evaluated on the same sequence-identity-reduced split as EPIESM-GA.

### 5.2 Evaluation protocol

We report AUC-ROC, PR-AUC, accuracy, F1, and Matthews correlation coefficient (MCC). AUCROC measures threshold-independent ranking quality (Fawcett, 2006); PR-AUC is useful under class imbalance; and MCC is included because it is more informative than accuracy when both classes matter (Chicco and Jurman, 2020). Results are reported as mean ± standard deviation across five seeds (42, 123, 256, 789, 1024), with resampled stratified splits, restarted GA optimization, and retrained classifiers. Following standard caution for repeated comparisons (Demšar, 2006), we additionally apply a paired Wilcoxon signed-rank test for GA-selected versus dense ESM-2 XGBoost results.

### 5.3 Responsible evaluation

We report non-accuracy criteria relevant to responsible foundation-model deployment: no PLM fine-tuning, downstream trainable parameters, selected feature dimension, and relative inference latency, following Green AI principles (Schwartz et al., 2020). Because scores may prioritize wet-lab validation, we also report the Brier score and expected calibration error using 10 equal-width bins.

## 6 Results

### 6.1 Main performance

Table 2 reports the main comparison.ESM-2 features outperform handcrafted and classical sequence representations, confirming the value of PLM-derived embeddings for epitope classification. EPIESM-GA achieves the best same-split performance, reaching 0.880 AUC and 0.740 MCC with only 420 GA-selected dimensions. It improves over both dense ESM-2 features (0.860 AUC, 0.710 MCC) and random 420-dimensional selection (0.849 AUC, 0.696 MCC), suggesting that evolutionary selection removes task-irrelevant embedding dimensions and improves downstream generalization. Published systems, especially BepiPred-3.0, provide useful literature context, but their reported scores are indicative because they were evaluated on different test sets.

**Table 1.**
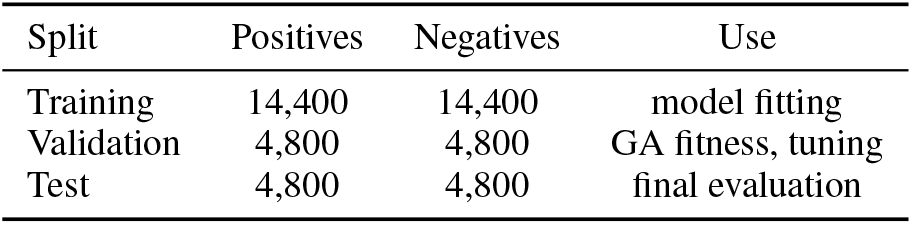
Balanced IEDB-derived split for linear B-cell epitope prediction. Counts follow the compact bench-mark setting used for the ESM-2 plus evolutionary selection experiments.

| Split | Positives | Negatives | Use |
| --- | --- | --- | --- |
| Training | 14,400 | 14,400 | model fitting |
| Validation | 4,800 | 4,800 | GA fitness, tuning |
| Test | 4,800 | 4,800 | final evaluation |

**Table 2.** Main predictive performance comparison. Published baselines marked with ^†^ are taken from their original papers and are therefore included only as indicative references, not as strictly controlled head-to-head comparisons. The ESM-2 variants in the lower block are evaluated on the same sequence-identity-reduced IEDB split used for EpiESM-GA.

| Method | Representation | AUC | PR-AUC | Acc. | F1<sup>a</sup> | MCC | Trainable |
| --- | --- | --- | --- | --- | --- | --- | --- |
| *Published baselines reported on their original evaluation settings* |  |  |  |  |  |  |  |
| BepiPred-2.0 (Jespersen et al., 2017) | Handcrafted | 0.740<sup>†</sup> | 0.690<sup>†</sup> | 0.690<sup>†</sup> | 0.680<sup>†</sup> | 0.580<sup>†</sup> | Small |
| EpitopeVec (Bahai et al., 2021) | PLM embedding | 0.820<sup>†</sup> | 0.760<sup>†</sup> | 0.760<sup>†</sup> | 0.730<sup>†</sup> | 0.650<sup>†</sup> | Small |
| LBCE-XGB (Zhu et al., 2023) | BERT embedding | 0.838<sup>†</sup> | 0.795<sup>†</sup> | 0.780<sup>†</sup> | 0.750<sup>†</sup> | 0.680<sup>†</sup> | XGBoost |
| BepiPred-3.0 (Clifford et al., 2022) | ESM-1b embedding | 0.857<sup>†</sup> | 0.810<sup>†</sup> | 0.790<sup>†</sup> | 0.758<sup>†</sup> | 0.700<sup>†</sup> | Medium |
| *Models evaluated on our sequence-identity-reduced IEDB split* |  |  |  |  |  |  |  |
| ESM-2 + XGBoost | Full ESM-2 | 0.860 | 0.830 | 0.800 | 0.770 | 0.710 | 1280-dim |
| ESM-2 + random selection | 420 ESM-2 dims | 0.849 | 0.815 | 0.792 | 0.761 | 0.696 | 420-dim |
| **EPIESM-GA** | **420 GA-selected ESM-2 dims** | **0.880** | **0.852** | **0.820** | **0.790** | **0.740** | **420-dim** |

### 6.2 Ablation: representational choices

Table 3 isolates the effect of feature representation. Composition-based features provide limited signal, while full ESM-2 mean pooling gives a large gain, indicating that contextual protein representations capture antigenic information beyond local residue counts. PCA and random subsampling compress ESM-2 but slightly reduce performance. In contrast, GA selection improves over the full 1280-dimensional representation, with gains of 0.028 AUC over PCA and 0.031 over random selection. This supports the view that the GA selects epitope-discriminative feature groups rather than merely reducing dimensionality.

**Table 3.** Feature creation and selection ablation (all using XGBoost). The primary representational gain comes from ESM-2 embeddings (Bahai et al., 2021; Clifford et al., 2022; Zhu et al., 2023). GA selection produces a more predictive compact representation than either PCA or random subsampling (Angaitkar et al., 2023). ^*a*^ F1 at threshold 0.5.

| Configuration | Dim. | AUC | F1<sup>a</sup> | MCC |
| --- | --- | --- | --- | --- |
| AA composition only | 20 | 0.711 | 0.641 | 0.402 |
| Dipeptide composition | 400 | 0.768 | 0.695 | 0.503 |
| ESM-2 mean pooled | 1280 | 0.860 | 0.770 | 0.710 |
| ESM-2 + PCA | 420 | 0.852 | 0.762 | 0.699 |
| ESM-2 + random | 420 | 0.849 | 0.761 | 0.696 |
| **ESM-2 + GA** | **420** | **0.880** | **0.790** | **0.740** |

### 6.3 Feature budget analysis

Table 4 studies the selected feature budget. A 128-dimensional subset loses substantial signal, whereas larger budgets increase latency with little added benefit. The 420-dimensional setting gives the best AUC and MCC while reducing classifier latency to 0.69× the dense 1280-dimensional baseline. The 640-dimensional setting is close, but the small single-split margin should be validated with cross-validation. Latency is measured after ESM-2 embeddings are cached.

**Table 4.** GA-selected feature budget sweep. Relative latency is normalized to 1280-dim dense classification (absolute: ≈42 ms per 512-peptide batch on A100, post-embedding-cache). ^*a*^ F1 at threshold 0.5.

| Dims | AUC | F1<sup>a</sup> | MCC | Rel. latency |
| --- | --- | --- | --- | --- |
| 128 | 0.833 | 0.741 | 0.671 | 0.41× |
| 256 | 0.861 | 0.772 | 0.712 | 0.57× |
| **420** | **0.880** | **0.790** | **0.740** | 0.69× |
| 640 | 0.878 | 0.788 | 0.737 | 0.82× |
| 1280 | 0.860 | 0.770 | 0.710 | 1.00× |

### 6.4 Classifier comparison

Table 5 compares downstream learners on the same 420 GA-selected ESM-2 features. XGBoost performs best, consistent with its ability to model nonlinear interactions. Random Forest remains competitive and provides feature-importance estimates, while the MLP is slightly weaker on this dataset. Logistic Regression lags behind tree-based models but confirms that the selected features retain a usable linear epitope signal.

**Table 5.** Downstream classifiers trained on the same 420 GA-selected ESM-2 features. ^*a*^ F1 at threshold 0.5.

| Classifier | AUC | PR-AUC | F1<sup>a</sup> | MCC |
| --- | --- | --- | --- | --- |
| Random Forest | 0.872 | 0.843 | 0.781 | 0.729 |
| XGBoost | **0.880** | **0.852** | **0.790** | **0.740** |
| MLP | 0.861 | 0.834 | 0.773 | 0.714 |
| Logistic Regression | 0.835 | 0.806 | 0.752 | 0.681 |

## 7 Discussion

Linear B-cell epitopes depend on interacting biochemical and contextual factors, including charge, hydrophilicity, solvent accessibility, flexibility, local motifs, and antigenic context (Potocnakova et al., 2016; Caoili, 2022). ESM-2 is useful because its embeddings encode local and long-range evolutionary dependencies, placing peptide residues in biochemical and structural neighborhoods even when only short fragments are available; this agrees with improvements reported for BepiPred-3.0 and other PLM-based epitope predictors (Clifford et al., 2022; Lin et al., 2023). GA selection is helpful because, unlike PCA or random subsampling, it directly optimizes held-out AUC and can retain synergistic dimensions, consistent with wrapper-based feature selection (Angaitkar et al., 2023); its 0.028 AUC gain over PCA supports this interpretation, pending confidence-interval validation. For deployment, EPIESM-GA reduces inputs from 1280 to 420 dimensions, lowers latency by about 31% (Table 4), and supports cached peptide screening; hardware-level energy and carbon profiling remains future work (Strubell et al., 2019; Schwartz et al., 2020; Patterson et al., 2021).

## 8 Conclusions

We proposed EPIESM-GA, a resource-efficient pipeline for linear B-cell epitope prediction that leverages ESM-2 as a frozen biological encoder, rather than a fine-tuned end-to-end model. EPIESM-GA compresses 1280-dimensional mean-pooled ESM-2 embeddings into 420 informative features using genetic algorithm-based selection and trains lightweight downstream classifiers to achieve 0.880 AUC-ROC and 0.740 MCC on a sequence-identity-reduced IEDB benchmark, with an inference latency reduction of roughly 31%. The ablation results show that ESM-2 contributes the dominant representational gain and that GA-based selection performs better than PCA and ran-dom compression at the same dimensionality. The proposed method does not require fine-tuning of PLM, works directly on amino acid sequences only, and outputs a small feature set that is suitable for compute-constrained screening workflows. This makes it relevant for peptide vaccine prioritization, pandemic preparedness, diagnostic antigen selection and antibody-discovery campaigns.

## 9 Reproducibility Details

We report the implementation details needed to reproduce the main experiments in Table 6. ESM-2 features were extracted using `esm2_t33_650M_UR50D` from `fair-esm` v2.0.0. The software stack included Python 3.10.12, PyTorch 2.1.0 with CUDA 11.8, scikit-learn 1.3.2, XGBoost 1.7.6, DEAP 1.4.1, NumPy 1.26.0, and Pandas 2.0.3. Embeddings were generated on one NVIDIA A100 40GB GPU with FP16 inference and batch size 64; GA optimization and classifier training used an Intel Xeon Gold 6338 CPU with 32 cores. Per seed, embedding extraction took about 45 minutes and GA optimization about 3 hours. Five seeds were used: 42, 123, 256, 789, and 1024. The IEDB data are available at https://www.iedb.org.

**Table 6.**
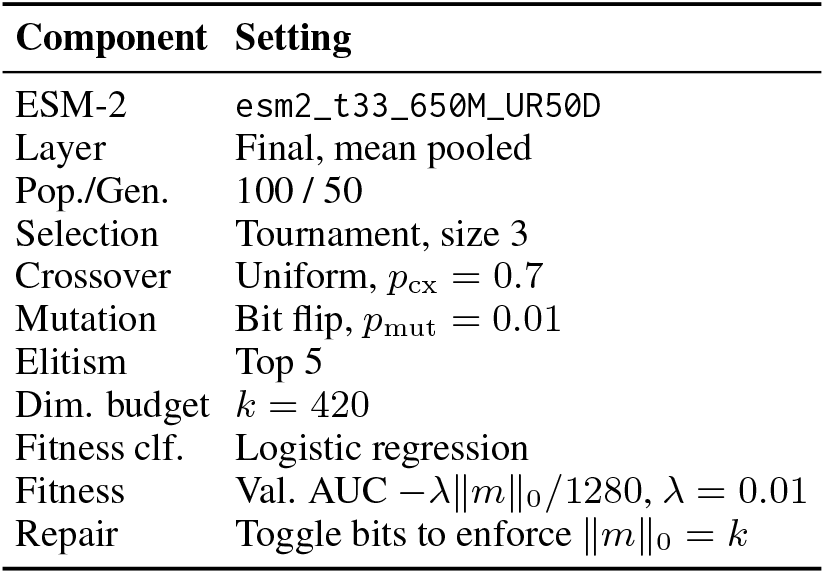
GA configuration for ESM-2 feature selection.

